# Characterization of SARS-CoV-2 variants B.1.617.1 (Kappa), B.1.617.2 (Delta) and B.1.618 on cell entry, host range, and sensitivity to convalescent plasma and ACE2 decoy receptor

**DOI:** 10.1101/2021.09.03.458829

**Authors:** Wenlin Ren, Xiaohui Ju, Mingli Gong, Jun Lan, Yanying Yu, Quanxin Long, Yu Zhang, Jin Zhong, Guocai Zhong, Xinquan Wang, Ailong Huang, Rong Zhang, Qiang Ding

**Affiliations:** Center for Infectious Disease Research, School of Medicine, Tsinghua University, Beijing 100084, China; School of Life Sciences, Tsinghua University, Beijing 100084, China; Key Laboratory of Molecular Biology on Infectious Diseases, Ministry of Education, Chongqing Medical University, Chongqing, China; Unit of Viral Hepatitis, CAS Key Laboratory of Molecular Virology and Immunology, Institut Pasteur of Shanghai, Chinese Academy of Sciences, Shanghai, China; Shenzhen Bay Laboratory, Shenzhen 518132, China; School of Chemical Biology and Biotechnology, Peking University Shenzhen Graduate School, Shenzhen 518055, China; Key Laboratory of Medical Molecular Virology (MOE/NHC/CAMS), School of Basic Medical Sciences, Shanghai Medical College, Biosafety Level 3 Laboratory, Fudan University, Shanghai 200032, China

**Keywords:** COVID-19, SARS-CoV-2, Delta variant, Kappa variant, B.1.618, host range, ACE2 decoy receptor

## Abstract

Recently, highly transmissible SARS-CoV-2 variants B.1.617.1 (Kappa), B.1.617.2 (Delta) and B.1.618 were identified in India with mutations within the spike proteins. The spike protein of Kappa contains four mutations E154K, L452R, E484Q and P681R, and Delta contains L452R, T478K and P681R, while B.1.618 spike harbors mutations Δ145-146 and E484K. However, it remains unknown whether these variants have altered in their entry efficiency, host tropism, and sensitivity to neutralizing antibodies as well as entry inhibitors. In this study, we found that Kappa, Delta or B.1.618 spike uses human ACE2 with no or slightly increased efficiency, while gains a significantly increased binding affinity with mouse, marmoset and koala ACE2 orthologs, which exhibits limited binding with WT spike. Furthermore, the P618R mutation leads to enhanced spike cleavage, which could facilitate viral entry. In addition, Kappa, Delta and B.1.618 exhibits a reduced sensitivity to neutralization by convalescent sera owning to the mutation of E484Q, T478K, Δ145-146 or E484K, but remains sensitive to entry inhibitors-ACE2-lg decoy receptor. Collectively, our study revealed that enhanced human and mouse ACE2 receptor engagement, increased spike cleavage and reduced sensitivity to neutralization antibodies of Kappa, Delta and B.1.618 may contribute to the rapid spread of these variants and expanded host range. Furthermore, our result also highlighted that ACE2-lg could be developed as broad-spectrum antiviral strategy against SARS-CoV-2 variants.

## INTRODUCTION

Since its emergence in late 2019, the severe acute respiratory syndrome coronavirus 2 (SARS-CoV-2) that causes the ongoing COVID-19 pandemic has evolved into several new viral variants of concern (VOC) and variants of interest (VOI)^1-4^. SARS-CoV-2 enters host cells by binding angiotensin-converting enzyme 2 (ACE2) in a species-dependent manner^2,5,6^. For example, murine, New World monkeys and koala ACE2 does not efficiently bind the SARS-CoV-2 spike protein, hindering viral entry into those species^7^. In addition, the spike is the target for vaccine and therapeutic antibodies^8,9^, and mutations in spike may potentially alter SARS-CoV-2 transmission, host tropism, pathogenicity as well as sensitivity to vaccine-elicited antibodies^10-12^. For example, D614G mutation, identified at the earlier stage of the pandemic, promotes spike binding to ACE2, leading to enhanced virus transmission^13,14^. Subsequently, the N501Y mutation found in the B.1.1.7, B.1.351 and B.1.1.28.1 spike has increased the binding affinity between the receptor-binding domain (RBD) and ACE2, increasing viral fitness and infectivity^1,15,16^; In addition, spike with the N501Y mutation has gained the ability to utilize mouse ACE2 as the receptor to infect mouse, expanding its host range^12^. In addition, K417N and E484K found in the B.1.351 variant contribute to evasion of neutralization by multiple monoclonal antibodies^17-19^. Thus, as the COVID-19 pandemic continues, it is critical to closely monitor the emergence of new variants, as well as their impact on viral transmission, pathogenesis, and vaccine and therapeutic efficacies.

Recently, the number of COVID-19 cases and deaths in India has risen steeply and the increased spread is associated with newly identified SARS-CoV-2 variants B.1.617 and B.1.618 with mutated spike proteins^20,21^. B.1.617.1 (Kappa), which carries E154K in the N-terminal domain (NTD) of spike, L452R and E484Q mutations in the RBD of spike, and P681R in proximity to furin cleavage site, has been designated as VOI by the World Health Organization (WHO) (https://www.who.int). B.1.617.2 (Delta) that carries L452R and T478K mutations in the RBD of spike, and P681R, has been designated as VOC (https://www.who.int). B.1.618 harbors Δ145-146 (deletion of 145^th^ and 146^th^ residues) and E484K mutation in the NTD and RBD, respectively. Indeed, some of the mutations in the Kappa, Delta and B.1.618 have been found in other variants separately. For example, the L452R mutation has been spotted in the B.1.427 and B.1.429 variants with enhanced transmissibility and reduced sensitivity to vaccine-elicited Abs^22,23^. T478K has been seen in Mexican variant B.1.1.519^24^. Also, the E484Q mutation is similar to the E484K found in the B.1.351, which exhibited reduce neutralization by convalescent sera or monoclonal antibodies^17,25-27^. For Kappa and Delta variants, this is the first time that L452R and E484Q (Kappa)/T478K (Delta) mutations are found to coexist together, and P681R is firstly emerged; for B.1.618, the combination of Δ145-146 in the NTD domainand E484K is firstly observed.

Here, we characterized the spike proteins of Kappa, Delta and B.1.618 on their ability to utilize different ACE2 orthologs for cell entry, and evaluated their sensitivity to convalescent sera and soluble ACE2-lg decoy receptor.

## RESULTS

### Characterization of cell entry driven by spike proteins of Kappa, Delta and B.1.618

The rapid spread of the new emerging variants could be caused by the increased ability to enter the cell, since the variants harbor mutations in the spike proteins (**Fig.1A**). To examine the biological impact of these mutations on cell entry, we produced pseudotyped virus particles containing a firefly luciferase reporter gene and expressing on their surface with the spike proteins of WT (D614G), Kappa, Delta and B.1.618 variants. HeLa cells expressing human ACE2 (HeLa-human ACE2) were then inoculated with these pseudoparticles and at 48h post-inoculation, the cells were lysed and the luciferase activity was monitored as a measure of virus entry (**Fig. 1B**). Compared to WT spike, Delta and B.1.618 spike proteins gained an increased ability to mediate viral entry into HeLa-human ACE2 cells, which are contributed by T478K (Delta), P681R (Delta), Δ145-146 (B.1.618) and E484K (B.1.618). Spike protein of Kappa exhibited comparable ability to mediate viral entry, even E484Q or P681R mutation in its spike could significantly promote viral entry individually.

**Figure 1.**
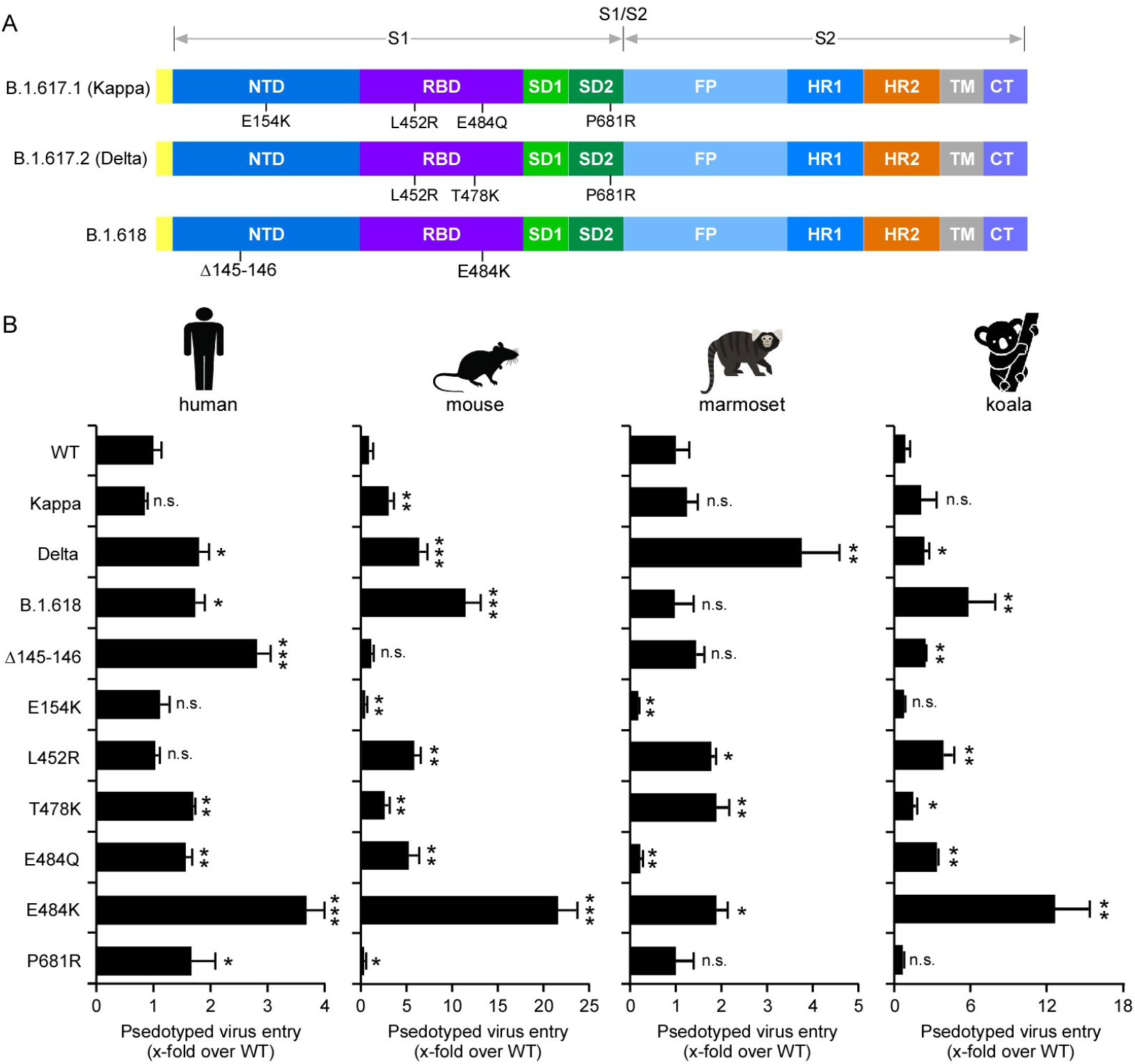
The spike proteins of variants driven viral entry into HeLa cells expressing human, mouse, marmoset or koala ACE2 orthologs. (A) Schematic overview of SARS-CoV-2 spike proteins of B.1.617.1 (Kappa), B.1.617.2 (Delta) and B.1.618, colored by domain. NTD, N-terminal domain; RBD, receptor binding domain; SD1, subdomain 1; SD2, subdomain 2; FP, fusion peptide; HR1, heptad repeat 1; HR2, heptad repeat 2; TM, transmembrane region; CT, cytoplasmic tail. (B) Cell entry of the virion pseudotyped with the WT, Kappa, Delta, B.1.618 spike proteins or individual mutations of these variants spike proteins were tested on HeLa cells expressing human, marmoset, or koala ACE2 orthologs. Luciferase activity was determined after 2 days of infection and data were normalized to the WT (D614G) of individual experiment. All infections were performed in triplicate, and the data are representative of three independent experiments (mean ± standard deviation). ns, no significance; *, P < 0.05, **, P < 0.01, ***, P < 0.001. Significance assessed by one-way ANOVA.

SARS-CoV-2 has a broad host range, and its spike could utilize a diverse range of ACE2 orthologs for cell entry^7,28^. However, we and others previously found that SARS-CoV-2 spike has a limited binding affinity with mouse, New World monkey or koala ACE2 and does not efficiently mediate virus entry into these species^7,28,29^. We thus sought to evaluate the abilities of variants’ spike proteins in utilization of these ACE2 proteins for cell entry. To this end, we produced virus pseudotyped with SARS-CoV-2 variant spike proteins with single or combination of mutations. The HeLa cells expressing mouse, marmoset (New World monkey), or koala ACE2 orthologs were then inoculated with pseudoparticles and luciferase activity was determined at 48h post-inoculation (**Fig. 1B**). Our results showed that spike proteins from Kappa, Delta and B.1.618 could significantly enhanced cell entry into HeLa-mouse ACE2 cells, as results of T478K (B.1.618), E484Q (Kappa) and E484K (Delta). Beside HeLa-mouse ACE2, Delta also exhibited significantly enhanced cell entry into HeLa-marmoset and HeLa-koala cells, which are contributed by L452R and T478K mutations. In contrast, Kappa did not increased cell entry into HeLa-marmoset or HeLa-koala cells, and B.1.618 only showed enhanced entry into HeLa-koala cells, which is attributable to E484K.

Taken together, our results demonstrated that the spike protein of Kappa, Delta or B.1.618 with distinct mutations have altered their ability in utilizing ACE2 orthologs for cell entry. Delta and B.1.618 variants gained an enhanced ability to use human ACE2 receptor for cell entry. Remarkably, Delta variant gained the function to utilize mouse, New World monkey or koala ACE2 orthologs, which cannot be engaged with WT virus, for cell entry, with potential to extend its host range to these species.

### The spike protein of Kappa, Delta and B.1.618 gained an increased binding affinity with human ACE2 and other othologs

As the Kappa, Delta and B.1.618 spike mediate increased cell entry efficiency, which could be caused by the increased binding affinity for ACE2. We employed a cell-based assay that uses flow cytometry to assess the binding of RBD of spike protein to human ACE2 (**Fig. S1A**). We cloned the cDNA of human ACE2 into a bicistronic lentiviral vector (pLVX-IRES-zsGreen1) that expresses the fluorescent protein zsGreen1 via an IRES element and can be used to monitor transduction efficiency. Next, WT or variants derived RBD-His (a purified fusion protein consisting of the RBD and a polyhistidine tag at the C-terminus) was incubated with HeLa cells transduced with the human ACE2. Binding of RBD-His to ACE2 was then quantified by flow cytometry (**Fig. S1 and Fig. 2A**). As shown, the binding efficiencies of the RBDs of Kappa (L452R+E484Q) (98.88%), Delta (L452R+T478K) (99.04%), B.1.618 (E484K) (98.76%), L452R (98.90%), and E484Q (98.48%) were higher than WT (89.6%), suggesting the RBD of variants bind human ACE2 with a higher affinity (**Fig. 2A**).

**Figure 2.**
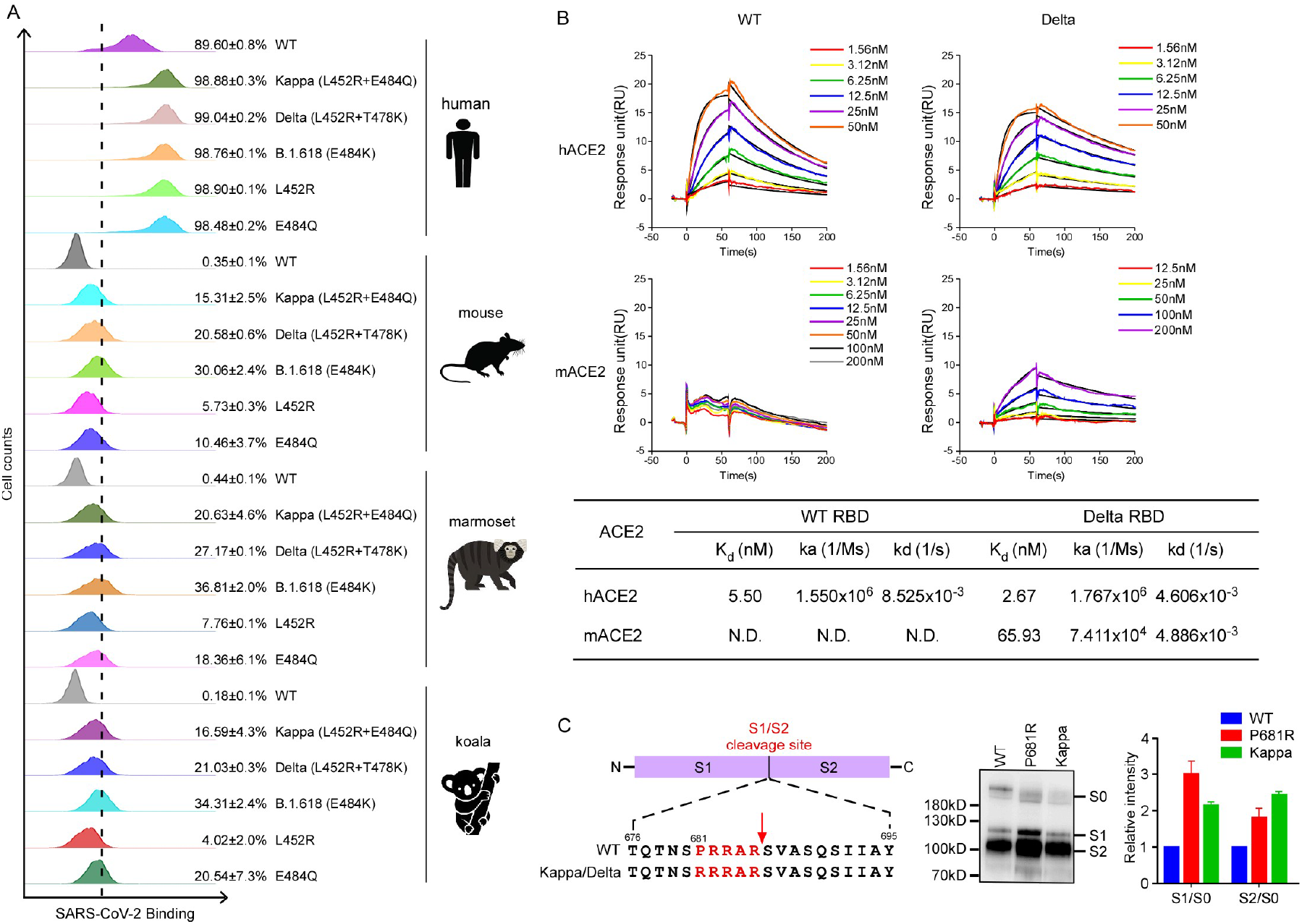
The binding of variants’ spike proteins with ACE2 orthologs and the spike protein processing. (A) HeLa-ACE2 cells were incubated with recombinant RBD-His proteins bearing mutations of Kappa, Delta and B.1.618 or individual mutation. The binding of RBD-His with cells were analyzed by flow cytometry. Values are expressed as the percent of cells positive for RBD-His among the ACE2-expressing cells (zsGreen1+ cells) and shown as the means ± SD from 3 biological replicates. This experiment was independently performed three times with similar results. (B) The binding kinetics of ACE2 proteins (human or mouse) with recombinant WT or Delta SARS-CoV-2 RBD were obtained using the BIAcore. ACE2 proteins were captured on the chip, and serial dilutions of RBD were then injected over the chip surface. Experiments were performed three times with similar result, and one set of representative data is displayed. (C) Immunoblot analysis of spike protein cleavage of pseudovirus of Kappa and P681R using polyclonal antibodies against spike. Full-length spike (S0), S1, and S2 protein are indicated. The ratio of the S1/S0 or S2/S0 was quantitatively analyzed using ImageJ software. This experiment is repeated twice independently, and data are normalized to the WT (D614G) of individual experiment.

To test whether the spike proteins of the variants have altered in binding with mouse, marmoset and koala ACE2 orthologs, we incubated the recombinant RBD-His of variants’ spike with HeLa cells expressing mouse, marmoset or koala ACE2, and the binding of RBD-His to ACE2 ortholog was quantified by flow cytometry (**Fig. S1 and Fig. 2A**).The WT RBD-His cannot bind with mouse, marmoset or koala ACE2 as previously reported^2,7,29^; In contrast, RBD-His of Kappa, Delta, and B.1.618 bind with mouse, marmoset and koala ACE2 with a varying affinity, suggesting that these variants have evolved to gain the function for binding with <u>non-human</u> ACE2 orthologs.

As Delta variant has now become the most dominant strain of the coronavirus circulating globally, we expressed and purified recombinant human and ACE2, as well as WT and Delta variant’s RBDs, and directly assayed the protein binding in vitro by surface plasmon resonance (SPR) analysis (**Fig. 2B**). The dissociation constant (Kd) for human ACE2 binding the WT RBD was 5.50 nM while that of Delta RBD was 2.67 nM, about 2-fold higher than WT RBD. As expected, the WT RBD cannot bind with mouse ACE2; strikingly, the Delta RBD could bind mouse ACE2 with Kd of 65.93 nM.

Taken together, our results demonstrated that the spike proteins of Kappa, Delta and B.1.618 have evolved to enhance their binding affinity with human ACE2. Remarkably, the spike proteins of these variants also gain the function to bind with mouse, marmoset and koala ACE2, with potential to extend SARS-CoV-2 host range.

### P681R mutation in Kappa and Delta variants with enhanced spike protein cleavage

SARS-CoV-2 spike harbors a multibasic furin cleavage site (residues 681–686; PRRARS) at the S1/S2 junction, and the proteolytic processing of the spike by furin and TMPRSS2 proteases is important for SARS-CoV-2 infection^5,30-33^. Kappa and Delta variants contain a P681R substitution (**Fig. 1A** and **2C**), potentially optimizing the furin cleavage site, which prompted us to examine the effect of the P681R substitution on furin cleavage. To do this, we produced MLV viral particles pseudotyped with WT spike, Kappa spike, or P681R spike. Viruses in the cell culture supernatants were harvested and concentrated for immunoblot analysis of spike protein cleavage by polyclonal antibody against spike protein (**Fig. 2C**). Interestingly, our data showed significantly increased cleavage of the full length spike protein (S0) into the S1 and S2 fragments in Kappa spike and P681R spike pseudotyped viruses compared with WT spike., The cleaved S1/S0 ratio was 2.1-(Kappa) or 3.0-(P681R) folds higher than WT, and the cleaved S2/S0 was 2.4-(Kappa) or 1.8 (P681R)-folds higher than WT. These results suggest that P618R substitution in the Kappa and Delta variants could enhance spike cleavage, and subsequently facilitate viral entry and transmission.

### Kappa, Delta and B.1.618 variants exhibited resistance to neutralization by convalescent serum, while remained sensitive to ACE2 based decoy receptor antiviral countermeasure

SARS-CoV-2 infection-elicited neutralizing antibodies target the spike protein, which is critical for protection from re-infection^34,35^. We hypothesized that mutations in the spike protein of the Kappa, Delta and B.1.618 variants might contribute to the evasion of neutralizing antibodies. Therefore, we sought to determine the sensitivity of these variants to neutralization by convalescent serum. We chose plasma from COVID-19 patients (**Table S1, S2 and S3**) and measured the neutralization activity of convalescent plasma against virions pseudotyped with single or combined mutations from Kappa, Delta and B.1.618 variants (**Fig. 4A, B and C**). To this end, we preincubated the serial-diluted convalescent sera with virion pseudotyped with spike proteins as describe above, and subsequently tested on HeLa-human ACE2 cells. Cell entry of pseudotyped virion in presence of convalescent plasma with varying concentrations was assessed 48 hours later by measurement of luciferase activities. The results showed that Kappa, Delta and B.1.618 exhibited 1.8-, 3.0- and 3.3-folds resistance to neutralization by convalescent sera, respectively, which is conferred by E484Q, L452R+E484Q, T478K, Δ145-146 and E484K (**Fig. 4A, B and C**).

Previous studies have shown that ACE2-lg (ACE2 fused with Fc recombinant protein) exhibited a potent antiviral effect against SARS-CoV-2 infection^28,36^. As the spike proteins of Kappa, Delta and B.1.618 exhibited enhanced ACE2 binding affinity, it could be more sensitive to the inhibition by ACE2 decoy receptor. To this end, we used an SARS-CoV-2 transcription and replication competent virus-like particle (trVLP) cell culture system, which recapitulates the entire viral life cycle in Caco-2-N cells^37^, to engineer the desired mutations in the spike proteins of Kappa, Delta and B.1.618 variants into an SARS-CoV-2 isolate Wuhan-Hu-1 with D614G (WT) backbone, and examined the sensitivity of trVLP of Kappa, Delta and B.1.618 to inhibition of ACE2-lg. Specifically, we inoculated the Caco-2-N cells with WT, Kappa, Delta or B.1.618 trVLP (moi=0.1) in the presence of ACE2-lg at varying concentrations. After 48h, the cells were collected and GFP expression was quantified as the proxy of virus infection by flow cytometry. ACE2-lg could potently inhibit WT, Kappa, Delta and B.1.618 trVLP infection with IC50 of 21.05 ng/ml, 9.90 ng/ml, 15.80ng/ml and 15.77 ng/ml, indicating that the Kappa, Delta and B.1.618 are still sensitive to inhibition by ACE2-lg (**Fig. 4A, B and C**). In summary, the Kappa, Delta and B.1.618 exhibited a reduced sensitivity to neutralization by convalescent serum, while remained sensitive to ACE2 decoy receptor antiviral countermeasure.

## DISCUSSION

The emergence of SARS-CoV-2 variants imposes challenges to control of the COVID-19 pandemic^1,38,39^. The recent surge in COVID-19 cases and mortalities in India is associated with new SARS-CoV-2 variants Kappa, Delta and B.1.618 with mutated spike proteins^21^. In this manuscript, we characterized the biological properties of these new variants, including the efficiency of entry into cells, the binding affinities with human ACE2, as well as other orthologs and the sensitivity to neutralization by convalescent plasma and recombinant ACE2-lg decoy receptor (**Fig. 5**).

We found that the Kappa variant has not increased in cell entry of HeLa-human ACE2 cells and slightly increased in cell entry of HeLa-mouse ACE2 cells. Delta variant has significantly increased in cell entry of HeLa-human ACE2, HeLa-mouse ACE2, HeLa-marmoset ACE2 and HeLa-koala ACE2 cells; consistently, Delta RBD binds human ACE2 with a higher affinity. Remarkably, Delta RBD gained the ability to bind with mouse, marmoset and koala ACE2 orthologs, which exhibited limited binding affinity with WT RBD. B.1.618 variant has dramatically increased in cell entry of HeLa-human ACE2 cells, HeLa-mouse ACE2, and HeLa-koala ACE2 cells, but not HeLa-marmoset ACE2 cells. Additionally, P681R mutation in the spike proteins of Kappa and Delta variants enhanced the spike processing. Recent studies have found that the Delta bearing P681R infection could form large size syncytia compared to other variants^40^, further indicating that the P681R mutation in the furin cleavage site could enhance viral fusogenicity. As furin cleavage site is critical for viral pathogenesis and transmission^32,41^, the Kappa and Delta bearing the optimized furin cleavage site have potentially evolved increased pathogenicity and transmissibility, which is urgently needed to be investigated. Remarkably, our results suggest that the Kappa, Delta and B.1.618 variants have extended their ACE2 orthologs usages into mouse, koala and New World monkeys (**Fig. 1B**), raising a potential risk of mice or other rodents becoming the reservoirs for SARS-CoV-2, and the virus could potentially spillback to humans as the mice are living closed to human (**Fig. 5**). Thus, we recommend that the host range should be closely monitored along the continued evolution of SARS-CoV-2 to prevent future zoonosis-associated outbreaks.

SARS-CoV-2 variants Kappa, Delta and B.1.618 exhibited reduced sensitivity to neutralization by polyclonal antibodies in the serum from individuals previously infected with SARS-CoV-2 (**Fig. 3A, B and C**). Our analysis suggest that the immune escape is mainly conferred by the E484Q, T484K, Δ145-146 and E484K mutations, these new variants must be further surveyed to avoid fast-spreading and raises alerts if it was considered to be soon variants contributing to accelerate the spread of the virus in human populations. Thus, our findings highlight the critical need for broad-spectrum neutralizing antibodies insensitive to substitutions arising in VOCs or VOIs. In addition, we further demonstrated that the ACE2 decoy receptor-based antiviral strategy was represented as an alternative countermeasures against the VOCs, as the VOCs, Kappa and B.1.618 exhibited increased binding with human ACE2^1,39^. Our results showed that recombinant ACE2-lg protein could inhibit Kappa, Delta and B.1.618 infection with efficacy of comparable or better than that of WT (**Fig. 4A-D and Fig. 5**), which is consistent with the results that Kappa, Delta and B.1.618 spike proteins exhibited increased binding affinity with human ACE2 (**Fig. 2B**). In addition, it has been demonstrated that human ACE2 peptidase activity and viral receptor activity could be uncoupled^42,43^, thus it is possible that the enzymatic inactivated ACE2 was developed as the antivirals in clinics to avoid the potential ACE2 side effected mediated by its enzymatic activity.

**Figure 3.**
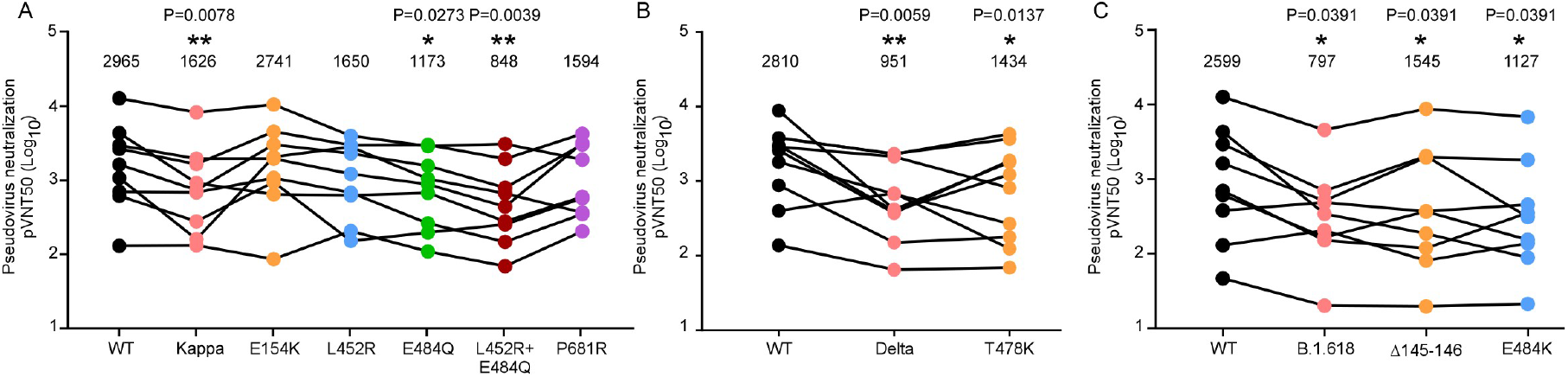
Reduced sensitivity of Kappa, Delta and B.1.618 to neutralization of convalescent sera. (A-C) MLV particles pseudotyped with the indicated spike proteins of Kappa, Delta and B.1.618 or mutations as indicated were preincubated with serially diluted convalescent sera, respectively. HeLa-ACE2 cells were incubated with these preincubated mixes and analyzed 48 h later by measuring luciferase activity to calculate the plasma dilution factor leading to 50% reduction in spike protein-driven cell entry (neutralizing titer 50, NT50). NT50 of each serum against each pseudovirion was presented and identical serum samples are connected with lines. Statistical significance of differences between WT and variant spike proteins was analyzed by two sided Friedman test with Dunn’s multiple comparison. *P < 0.05; **P < 0.01. pVNT50 of each sample is tested by two repeat.

**Figure 4.**
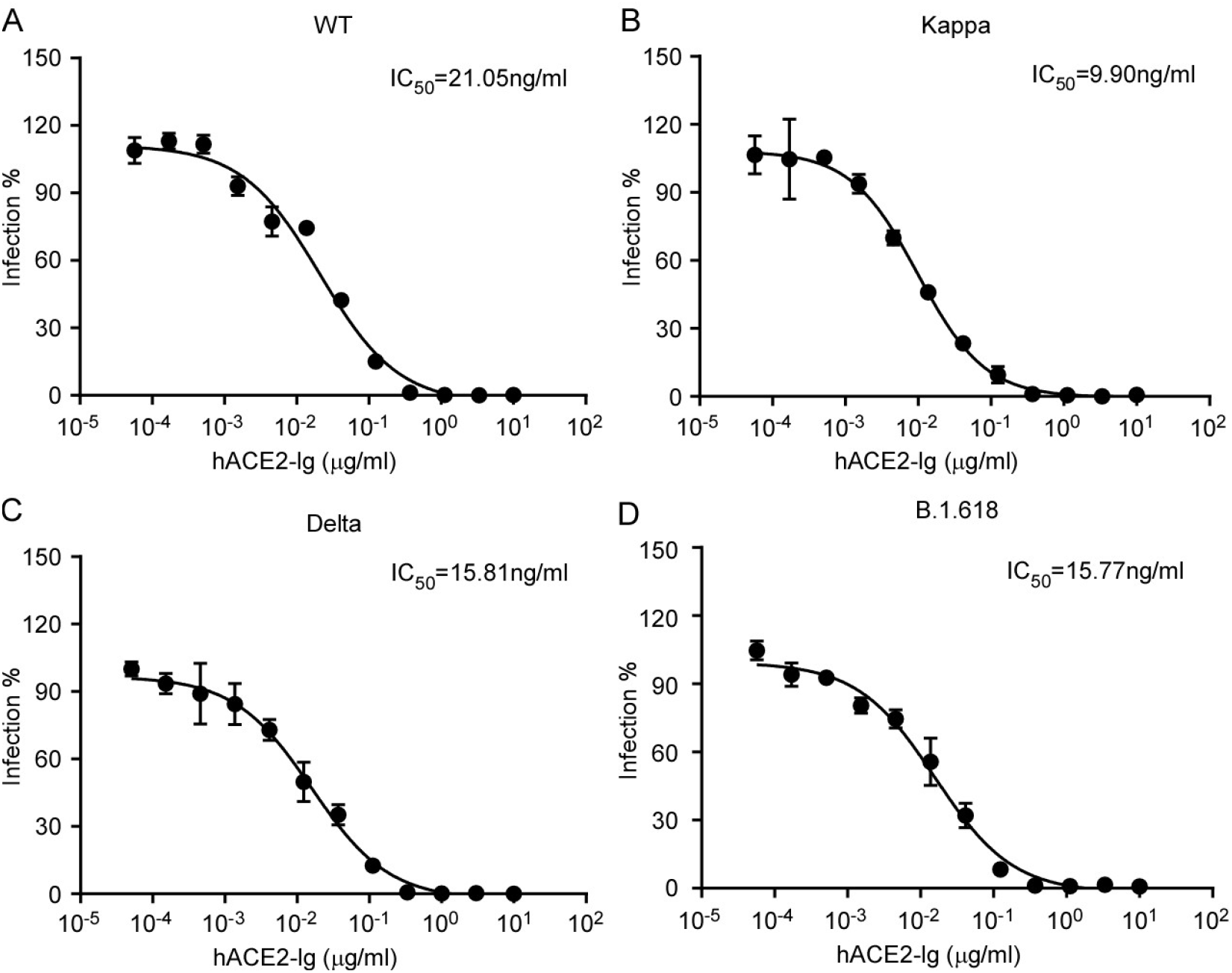
Inhibition of WT, Kappa, Delta and B.1.618 by recombinant ACE2-lg decoy receptor. Recombinant ACE2-Ig was diluted at the indicated concentrations. Viral entry was determined by assessing Luc activity 48 hours post infection of WT (A), Kappa (B), Delta (C) and B.1.618 (D) trVLP. The dilution factors leading to 50% reduction of pseudotyped virion entry was calculated as the IC_50_using GraphPad Prism software. Data shown are representative of three independent experiments with similar results, and data points represent mean ± SD in triplicate.

**Figure 5.**
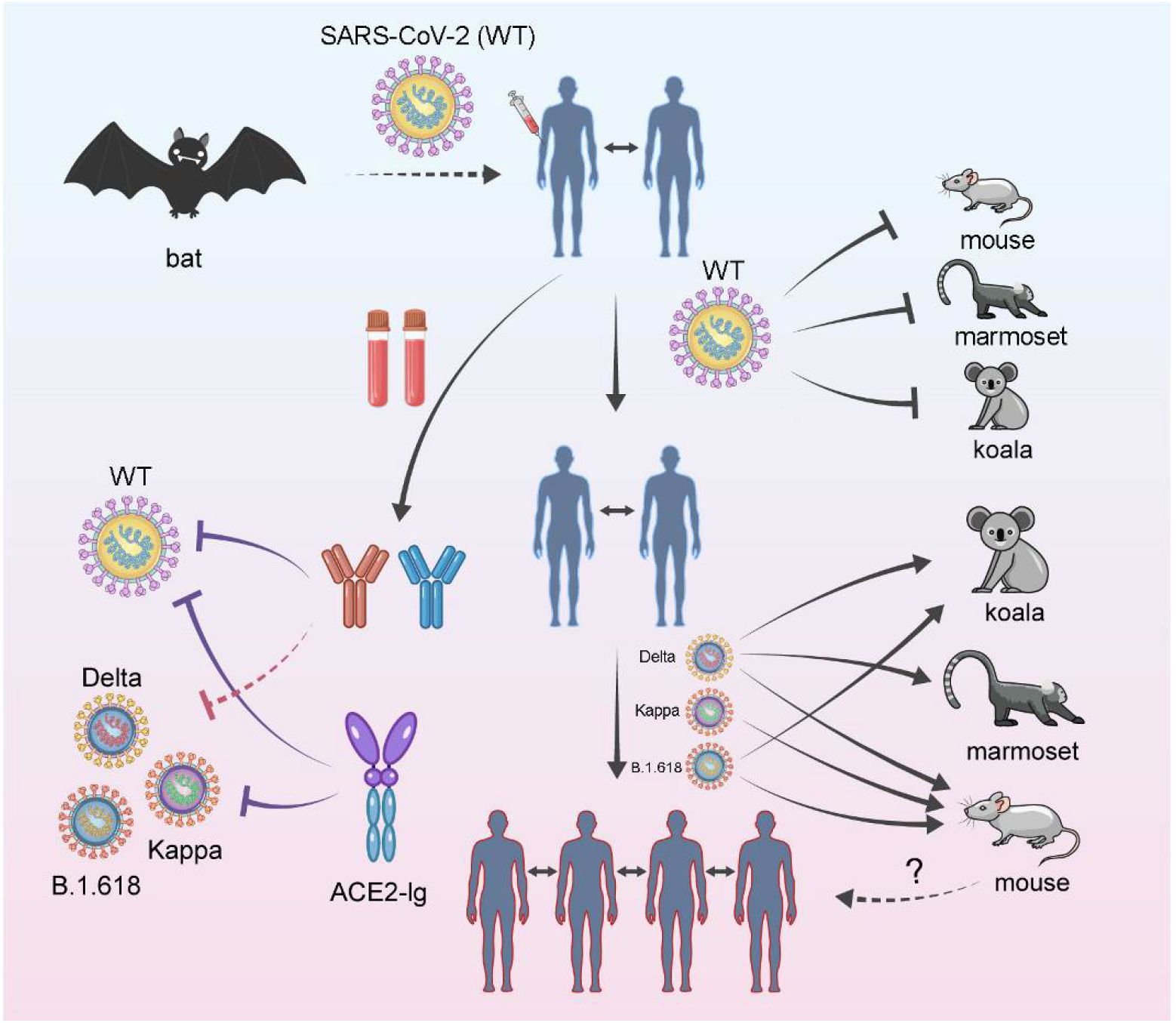
Schematic summary of SARS-CoV-2 variants Kappa, Delta and B.1.618 on cell entry, ACE2 orthologs utilization, and their sensitivities to convalescent plasma and ACE2 decoy receptor. Bats are considered as the natural zoonotic reservoir for SARS-CoV-2, which is the causative agent for COVID-19. SARS-CoV-2 uses ACE2 as the receptor to enter host cells in a species-dependent manner. ACE2 orthologs of mouse, marmoset or koala cannot bind with SARS-CoV-2 spike protein to mediate virus entry, therefore, these species are not permissive to SARS-CoV-2 infection. Several viral variants have been emerged, such as Kappa, Delta and B.1.618 harboring mutations in the RBD of spike proteins, and these variants have evolved increased binding affinity with human ACE2, as well as ACE2 orthologs of mouse, marmoset or koala, which potentially increased transmission in humans, extended their host range, and posed the risk of zoonotic transmission of virus into humans as mouse have closed contact with humans. In addition, these variants exhibited reduced sensitivities to convalescent plasma from the recovered patients infected by SARS-CoV-2 (WT), and still sensitive to ACE2-lg decoy receptor antiviral measurement.

New variants of concern will continue to emerge as the COVID-19 pandemic persists, which highlight the importance of genomic surveillance for the early identification of future variants. The potential of variants to escape naturally induced and vaccine elicited immunity makes the development of next-generation vaccines that elicit broadly neutralizing activity against current and future variants a priority^22,25,40^. In addition, the suppression of viral replication with both public health measures and the equitable distribution of vaccines, increasing the proportion of the population immunized with current safe and effective authorized vaccines, is critical to minimize the risk of emergence of new variants. Also the development of broad-spectrum antivirals, especially against diverse SARS-CoV-2 variants, is therefore of continued significance.

## Acknowledgements

We thank Dr. Jenna M. Gaska for suggestions and revision of the manuscript. We are grateful to other members of the Ding lab for critical discussions and comments on the manuscript.

This work was supported by the National Natural Science Foundation of China (32070153 to QD), Beijing Municipal Natural Science Foundation (M21001 to QD), and Start-up Foundation of Tsinghua University (53332101319).

## Materials and methods

### Cell culture

HEK293T (American Tissue Culture Collection, ATCC, Manassas, VA, CRL-3216), Vero E6 (Cell Bank of the Chinese Academy of Sciences, Shanghai, China) and A549 (ATCC) cells were maintained in Dulbecco’s modified Eagle medium (DMEM) (Gibco, NY, USA) supplemented with 10% (vol/vol) fetal bovine serum (FBS), 10mM HEPES, 1mM sodium pyruvate, 1×non-essential amino acids, and 50 IU/ml penicillin/streptomycin in a humidified 5% (vol/vol) CO2 incubator at 37°C. Cells were tested routinely and found to be free of mycoplasma contamination.

### Plasmids

The cDNAs encoding the ACE2 orthologs were synthesized by GenScript and cloned into the pLVX-IRES-zsGreen1 vector (Catalog No. 632187, Clontech Laboratories, Inc) with a C-terminal FLAG tag. ACE2 mutants were generated by Quikchange (Stratagene) site-directed mutagenesis. All constructs were verified by Sanger sequencing.

### Lentivirus production

Vesicular stomatitis virus G protein (VSV-G) pseudotyped lentiviruses expressing ACE2 orthologs tagged with FLAG at the C-terminus were produced by transient co-transfection of the third-generation packaging plasmids pMD2G (Addgene #12259) and psPAX2 (Addgene #12260) and the transfer vector with VigoFect DNA transfection reagent (Vigorous) into HEK293T cells. The medium was changed 12 h post transfection. Supernatants were collected at 24 and 48h after transfection, pooled, passed through a 0.45-µm filter, and frozen at -80°C.

### Surface ACE2 binding with RBD-His assay

HeLa cells were transduced with lentiviruses expressing the ACE2 variants for 48 h. The cells were collected with TrypLE (Thermo #12605010) and washed twice with cold PBS. Live cells were incubated with the recombinant proteins RBD-His with mutations (Sino Biological Cat. #40592-V08B; 40592-V08H88, 40592-V08H90, 40592-V08H84, 40592-V08H28, 40592-V08H81, 1μg/ml) at 4°C for 30 min. After washing, cells were stained with Anti-His-PE (clone: GG11-8F3.5.1; Miltenyi Biotec; Cat. #130-120-787) for 30 min at 4°C. Cells were then washed twice and subjected to flow cytometry analysis (Thermo, Attune™ NxT). Binding efficiencies are expressed as the percent of cells positive for RBD-His among the zsGreen positive cells (ACE2 expressing cells).

### Surface plasmon resonance analysis

The WT or Delta SARS-CoV-2 RBD (residues Arg319–Phe541) and the N-terminal peptidase domain of human or mouse ACE2 (residues Ser19–Asp615) were expressed using the Bac-to-Bac baculovirus system (Invitrogen) as described previously^44^. ACE2 was immobilized on a CM5 chip (GE Healthcare) to a level of around 500 response units using a Biacore T200 (GE Healthcare) and a running buffer (10 mM HEPES pH 7.2, 150 mM NaCl and 0.05% Tween-20). Serial dilutions of the SARS-CoV-2 RBD were flowed through with a concentration ranging from 400 to 12.5 nM. The resulting data were fit to a 1:1 binding model using Biacore Evaluation Software (GE Healthcare).

### Production of SARS-CoV-2 pseudotyped virus, determination of viral entry efficiency and analysis of spike protein cleavage

Pseudoviruses were produced in HEK293T cells by co-transfecting the retroviral vector pTG-MLV-Fluc, pTG-MLV-Gag-pol, and pcDNA3.1 expressing SARS-CoV-2 spike gene or VSV-G (pMD2.G (Addgene #12259)) using VigoFect (Vigorous Biotechnology). At 48 h post transfection, the cell culture medium was collected for centrifugation at 3500 rpm for 10 min, and then the supernatant was subsequently aliquoted and stored at -80°C for further use. Virus entry was assessed by transduction of pseudoviruses in cells expressing ACE2 ortholog or mutants in 48-well plates. After 48 h, intracellular luciferase activity was determined using the Luciferase Assay System (Promega, Cat. #E1500) according to the manufacturer’s instructions. Luminescence was recorded on a GloMax® Discover System (Promega). To analysis of the spike protein cleavage, the concentrated pseudoviruses were produced by ultracentrifugation at 100,000g for 2 h over a 20% sucrose cushion. Western Blot detection of SARS-CoV-2 Spike protein was performed using a polyclonal Spike antibody (Sino Biological Cat. # 40589-V08B1).

### Human convalescent serum and neutralization of pseudotyped virion particles

We obtained convalescent serum from COVID-19 patients (**Table S1, S2** and **S3**) more than one month after documented SARS-CoV-2 infection in the spring of 2020. Each plasma sample was heat-inactivated (56°C, 30 min) and then assayed for neutralization against WT or mink-variant pseudoviruses. For neutralization experiments, S protein bearing pseudotyped virion particles were pre-incubated for 30 min at 37°C with diluted plasma samples obtained from convalescent COVID-19 patients, before the mixtures were inoculated onto HeLa-ACE2 cells. Transduction efficiency was determined at 48 h post inoculation. This study was approved by the Institution Review Board of Tsinghua University (20210040).

### Recombinant ACE2-lg protein expression and purification

ACE2-lg, a recombinant Fc fusion protein of soluble human ACE2 (residues Gln18-Ser740) was expressed in 293F cells and purified using protein A affinity chromatography as described in our previous study^28^.

### Production of SARS-CoV-2 trVLP

The desired mutations in the spike proteins of Kappa, Delta and B.1.618 variants into an SARS-CoV-2 isolate Wuhan-Hu-1 with D614G (WT) backbone, and the trVLP were generated as previously described^37^.

### Statistical analysis

One-way analysis of variance (ANOVA) with Tukey’s honestly significant difference (HSD) test was used to test for statistical significance of differences between the different group parameters. *P* values of less than 0.05 were considered statistically significant.

## Supplemental Figures and Figure legends

**Supplemental Figure 1.**
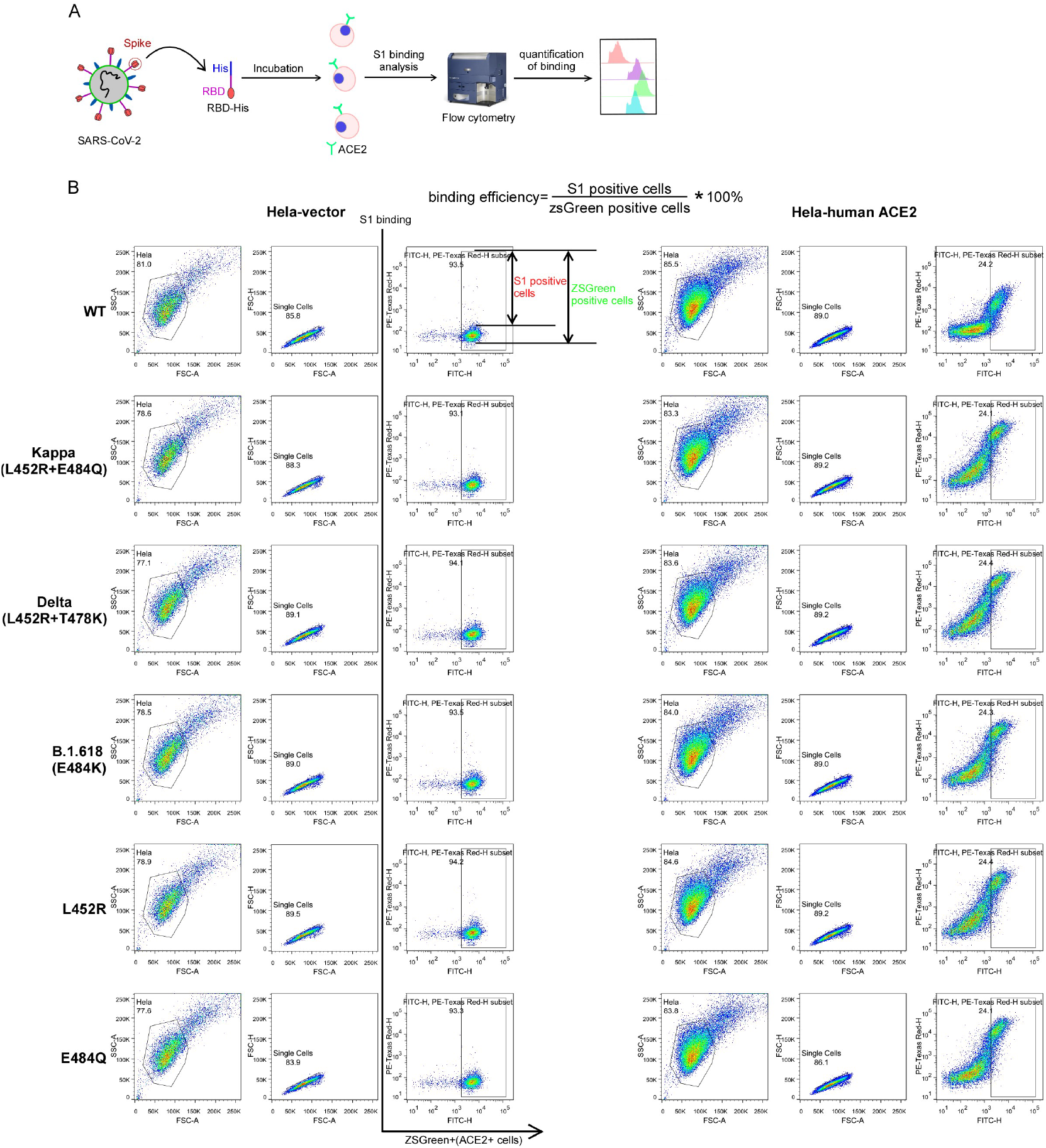
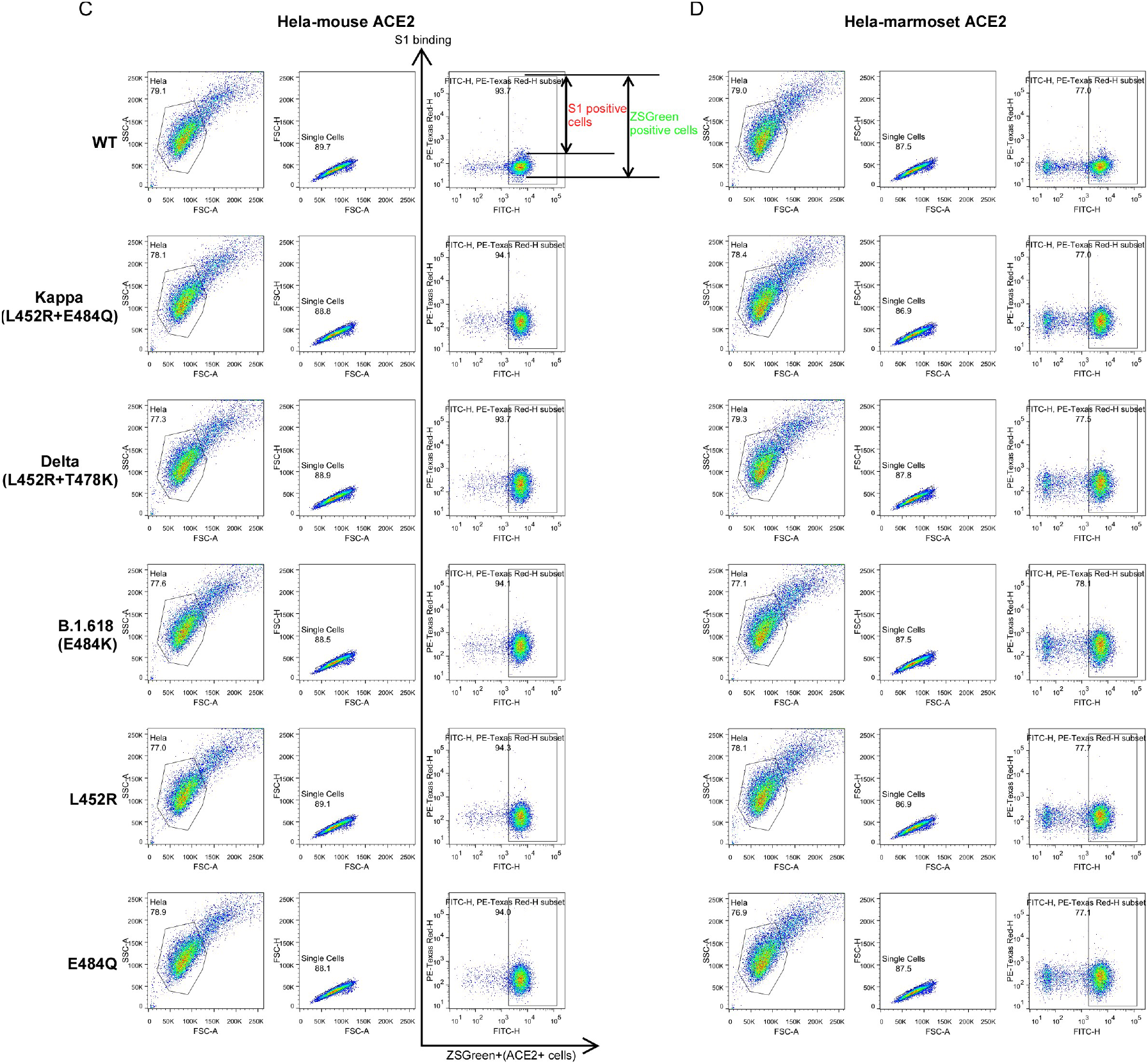
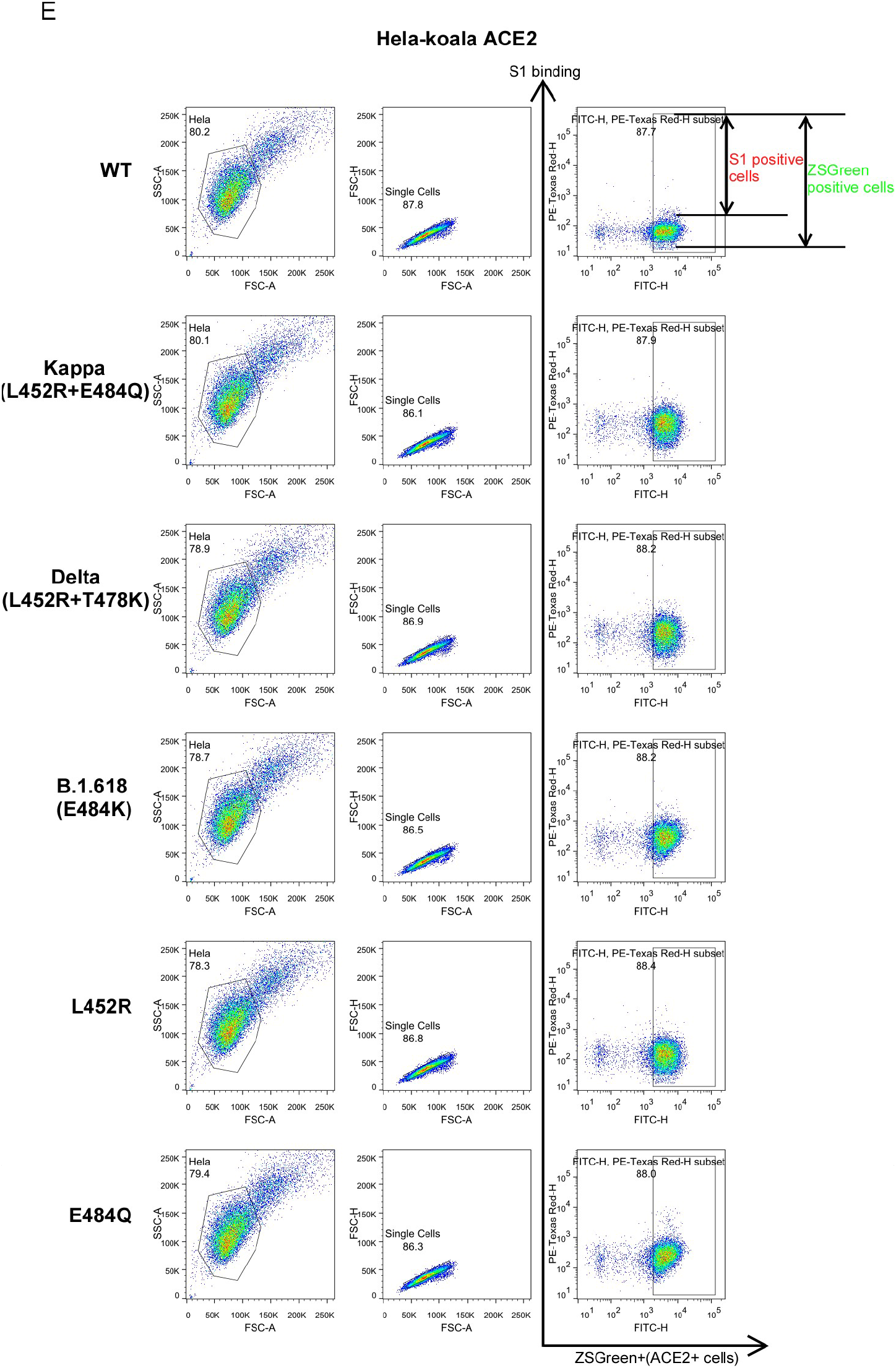
Gating strategy for determination of the binding efficiency of ACE2 variants with SARS-CoV-2 RBD-His protein. (A) Schematic of testing the efficiency of ACE2 binding with recombinant viral RBD-His protein. (B-E) Main cell population was identified and gated on Forward and Side Scatter. Single cells were further gated on FSC-A and FSC-H. The gated cells were plotted by FITC-A (zsGreen, as the ACE2 expressing population) and APC-A (RBD-His bound population). The FITC-A positive cell population was plotted to show the RBD-His positive population as Fig 2B. The binding efficiency was defined as the percent of RBD-His binding cells among the zsGreen positive cells. Shown are FACS plots representative of those that have been used for the calculations of binding efficiencies of ACE2 variants with RBD-His. This experiment was independently repeated three times with similar results.

## Supplemental Tables

**Table S1.** Summary of sera information of COVID-19 patients for evaluation of Kappa sensitivity.

Kappa sensitivity
| Characteristics | Patients (n=9) |
| --- | --- |
| **Age (median, range)** | 50 (15-69) |
| **Sex** |  |
| Male (%) | 2 (22.2) |
| Female (%) | 7 (77.8) |
| **Disease severity** |  |
| Severe | 4 (44.4) |
| Non-severe | 5 (55.6) |
| **Comorbidities, n (%)** |  |
| Fever | 5 (55.6) |
| Fatigue | 3 (33.3) |
| Dry cough | 1 (11.1) |
| Dyspnea | 4 (44.4) |
| Expectoration | 6 (66.7) |
| Nausea | 2 (22.2) |
| Dizziness | 1 (11.1) |
| Chills | 1 (11.1) |
| Chest stuffiness | 4 (44.4) |

**Table S2.** Summary of sera information of COVID-19 patients for evaluation of Delta sensitivity.

| Characteristics | Patients (n=10) |
| --- | --- |
| **Age (median, range)** | 52 (15-69) |
| **Sex** |  |
| Male (%) | 2 (20.0) |
| Female (%) | 8 (80.0) |
| **Disease severity** |  |
| Severe | 4 (40.0) |
| Non-severe | 6 (60.0) |
| **Comorbidities, n (%)** |  |
| Fever | 6 (60.0) |
| Fatigue | 4 (40.0) |
| Dry cough | 1 (10.0) |
| Dyspnea | 4 (40.0) |
| Expectoration | 6 (60.0) |
| Nausea | 2 (20.0) |
| Dizziness | 2 (20.0) |
| Chills | 1 (10.0) |
| Chest stuffiness | 5 (50.0) |

**Table S3.** Summary of sera information of COVID-19 patients for evaluation of B.1.618 sensitivity.

| Characteristics | Patients (n=9) |
| --- | --- |
| **Age (median, range)** | 49 (15-69) |
| **Sex** |  |
| Male (%) | 2 (22.2) |
| Female (%) | 7 (77.8) |
| **Disease severity** |  |
| Severe | 2 (22.2) |
| Non-severe | 7 (77.8) |
| **Comorbidities, n (%)** |  |
| Fever | 5 (55.6) |
| Fatigue | 3 (33.3) |
| Dry cough | 1 (11.1) |
| Dyspnea | 2 (22.2) |
| Expectoration | 4 (44.4) |
| Nausea | 1 (11.1) |
| Dizziness | 2 (22.2) |
| Chills | 1 (11.1) |
| Chest stuffiness | 4 (44.4) |

## Notes

### Competing Interest Statement

The authors have declared no competing interest.

